# SnRK1 acts upstream of *PRODUCTION OF ANTHOCYANIN PIGMENT1*, contributing to fine-tuning flavonoid biosynthesis during acclimation

**DOI:** 10.1101/2024.06.13.598896

**Authors:** Josephine Dieckmann, Ralf Stracke, Andreas S. Richter

## Abstract

In land plants, one of the processes vital to cope with environmental changes is the accumulation of photoprotective flavonoids such as flavonols and anthocyanins. The inactivation of SUCROSE NON-FERMENTING1 RELATED PROTEIN KINASE1 (SnRK1), which acts in a chloroplast-derived sugar signalling pathway, permits the activation of flavonoid biosynthesis in high-light. The present study provides genetic evidence that SnRK1 acts upstream of *PRODUCTION OF ANTHOCYANIN PIGMENT1 (PAP1)*, encoding a crucial transcription factor that activates the anthocyanin branch of flavonoid biosynthesis during high-light acclimation. A time-resolved expression analysis indicates a two-step suppression of MYB LIKE2 (MYBL2), a repressor of anthocyanin production, involving SnRK1 inactivation for stable anthocyanin accumulation during prolonged high-light exposure. Furthermore, overexpression of *PAP1* resulted in the marked suppression of *MYB11, MYB12* and *MYB111* and *FLAVONOL SYNTHASE1*, initiating the flavonol branch of the pathway. Analysis of a flavonoid-deficient *CHALCONE SYNTHASE* mutant overexpressing *PAP1* suggests that regulation of flavonoid biosynthesis is independent of flavonoid intermediates and end products but PAP1-dependent. It is proposed that PAP1 suppresses the flavonol branch by an as yet unknown mechanism, thereby promoting the consumption of carbon building blocks for anthocyanin production to permit the fine-tuning of the pathway.

## Introduction

Plants are sessile organisms unable to evade stressful environmental conditions. Among others, changes in incident light intensities put a thread on the cellular homeostasis, growth and, in its extremes, even survival of phototrophic organisms. Hence, multifactorial acclimation processes evolved to cope with the ever-changing environment. Among others, the accumulation of different flavonoids, such as flavonols and anthocyanins, is commonly observed when plants face challenging growth conditions. Among other eco-physiological functions, they are proposed to shield biomolecules from excessive amounts of (UV) light and to participate in the quenching of reactive oxygen species (Emiliani et al., 2013, Nakabayashi et al., 2014, Agati et al., 2020).

Anthocyanins are synthesised through the activities of enzymes that provide the precursors for all flavonoids, such as CHALCONE SYNTHASE (CHS) or CHALCONE ISOMERASE (CHI). The pathway then diverges into branches such as flavonol (glycosyl) and anthocyanin biosynthesis. FLAVONOL SYNTHASE1 (FLS1) catalyses the first step of the flavonol branch. The two products of the FLS1 reaction in *Arabidopsis thaliana*, kaempferol (K) and quercetin (Q), are regiospecifically modified by UDP-glycosyl transferases, which add glucose or rhamnose moieties (e.g., flavonol 3-O-rhamnosyltransferase UGT78D1) or methyltransferases, which methylate the flavonols (e.g., flavonol 3’O-Methyltransferase 1 (OMT1)). In a second major branch, DIHYDROFLAVONOL REDUCTASE (DFR), LEUCOCYANIDIN DIOXYGENASE (LDOX) (synonym ANTHOCYANINSYNTHASE (ANS)) and specific acyl-, methyl- and glycosyltransferases, like anthocyanin 5-O-glucoside malonyltransferase (5MAT), and sinapoyl glucose transferase (SCPL10), are involved in the final steps of anthocyanin production (Saito et al., 2013).

Given its complexity and number of branches, flavonoid biosynthesis is under spatio-temporal control of several transcription factors that manipulate the pathway for producing specific end-products in a given organ, tissue, or developmental state. For instance, Subgroup (SG) 7 MYB-transcription factors MYB11, MYB12 and MYB111 act as positive regulators of *CHS, CHI, FLS1* and glucosyltransferases coding genes in a tissue-specific context but also partially redundant manner (Stracke et al., 2007, Dubos et al., 2010, Stracke et al., 2010b). In Arabidopsis, activation of the anthocyanin branch involves SG6 MYBs encoded by *MYB75, MYB90, MYB113 and MYB114* (Borevitz et al., 2000; Gonzalez et al. 2008). SG6 transcription factors consist of a conserved MYB DNA-binding domain composed of sequence repeats (R) and a domain essential for interacting with basic helix-loop-helix (bHLH) transcription factors. At least one SG6 MYB is encoded in the genome of different anthocyanin-accumulating species, such as Arabidopsis, grape, petunia or snapdragon (Li, 2014, LaFountain and Yuan, 2021). Together with a WD40 protein (in Arabidopsis TRANSPARENT TESTA GLABRA1 (TTG1)), MYB and bHLH form a ternary regulative complex known as <u>M</u>YB-<u>b</u>HLH-<u>W</u>D40 (MBW) complex.

The MBW complex predominantly stimulates the expression of gene products involved in (pro-) anthocyanin biosynthesis, such as DFR and LDOX, but also transcripts for enzymes involved in early steps of the pathway, for example, CHS and CHI (Tohge et al., 2005, Nakabayashi et al., 2014). Although the other three Arabidopsis SG6 MYB factors act positively on flavonoid biosynthesis genes, PRODUCTION OF ANTHOCYANIN PIGMENT1 (PAP1, encoded by *MYB75*) plays a central role in the activation of anthocyanin biosynthesis (Borevitz et al., 2000, Zheng et al., 2019). In contrast to the activators, several MYB-type repressors of structural flavonoid biosynthesis genes have been identified. Based on phylogenetic relationships and specific sequence motifs, the repressors are grouped into the R3-MYB CPC (CAPRICE) or the SG4 R2R3-MYBs type. The repressive MYBs act either as inhibitors of the expression of flavonoid biosynthesis genes through an EAR-repression motif or by repressing the activity of the MBW complex, most likely through competition with SG6 MYB components for binding to the MBW complex (LaFountain and Yuan, 2021). Besides a general negative function on flavonoid biosynthesis genes, some repressive MYBs are also induced by the MBW complex, suggesting that they act to fine-tune the expression and prevent the overproduction of flavonoids via feedback regulation (Li, 2014, LaFountain and Yuan, 2021). One of the SG4-derived R3-MYBs, MYB-like 2 (MYBL2), concurrently interacts with PAP1 for binding to the MBW complex, thereby repressing its activity (Dubos et al., 2008, Matsui et al., 2008, Wang et al., 2016). The activation of flavonoid biosynthesis is intertwined with light-signalling pathways through ELONGATED HYPOCOTYL5 (HY5) and its cofactors BBOX-PROTEIN (BBX) BBX20, BBX21, BBX22, which stimulate the expression of flavonoid biosynthesis enzymes and transcription factors, such as *PAP1* or *MYB12*, through binding to the respective promoters (e.g., Stracke et al., 2010a, Shin et al., 2013, Bursch et al., 2020). Deficiency of these factors results in lower anthocyanin content in high-light and after exposure to monochromatic light (Kleine et al., 2007, Stracke et al., 2010a, Bursch et al., 2020). It was also revealed that HY5 binding in two Z-boxes of the promoter of *MYBL2* and HY5-dependent suppression of *MYBL2* contribute to the activation of flavonoid biosynthesis (Wang et al., 2016).

Accumulation of anthocyanins is observed under various conditions, and multiple stimuli and signaling pathways involving the chloroplasts to activate anthocyanin biosynthesis have been identified (e.g., LaFountain and Yuan, 2021, Araguirang and Richter, 2022, Richter et al., 2023). Increased light intensity stimulates photosynthesis and results in increased cellular sugar content. Besides providing carbon (C) skeletons for anabolic and catabolic processes, intermediates and end-products of sugar biosynthesis pathway(s) also function as signals to adjust gene expression, metabolic reactions and growth (Li et al., 2021). One decisive component that senses plant cells’ cellular energy/ carbohydrate status is the SUCROSE NON-FERMENTING RELATED KINASE1 (SnRK1, Baena-Gonzalez et al., 2007). In *Arabidopsis*, SnRK1 consists of two catalytic α subunits (encoded by *KIN10* and *KIN11*), β and βγ subunits acting as scaffold and regulatory subunits, respectively (Crepin and Rolland, 2019). In plants, SnRK1 is proposed to be inactivated by regulatory sugar phosphates, permitting biosynthesis of C- and energy-demanding biomolecules when carbohydrate and energy levels are high (Baena-Gonzalez and Lunn, 2020). SnRK1 inactivation was shown to be a prerequisite for the activation of anthocyanin biosynthesis during high-light acclimation (Zirngibl et al., 2023). SnRK1 acts on anthocyanin accumulation through a post-translational mechanism involving phosphorylation, reallocation between cytosol and nucleus, and degradation of MBW-complex components, such as PAP1 (Broucke et al., 2023). However, a surplus of functional SnRK1 through the overexpression of the catalytic subunit *KIN10* resulted in the suppression of *PAP1* and anthocyanin biosynthesis gene expression (Zirngibl et al., 2023) and a more robust expression of *MYBL2* in protoplast assays (Ramon et al., 2019, Broucke et al., 2023). Hence, an impact of SnRK1 on the expression of pathway genes and regulators is indicated, but the factors involved downstream of SnRK1 remain to be identified.

The flavan scaffold almost exclusively consists of carbon, and the most abundant end-products of the anthocyanin branch accumulate as (poly-)glycosylated molecules (up to four carbohydrates per anthocyanin). Hence, biosynthesis of the different flavonoids is highly energy- and carbon-demanding (Hernandez and Van Breusegem, 2010, Saito et al., 2013) and must be tightly controlled and adjusted to the cellular energy/carbon status and cellular demands under environmental stress. In this context, activation of the different branches of flavonoid biosynthesis on all layers of gene expression to ensure a demand-driven distribution of the available precursors for the end products has to be controlled. How this is achieved in acclimation-relevant conditions is not resolved.

Here, we show that diminished activation of the pathway in the presence of active SnRK1 can be bypassed by artificial expression of *PAP1*, suggesting that SnRK1 acts upstream of *PAP1* and analysis of flavonol and anthocyanin content indicates a more predominant effect of SnRK1 on the anthocyanin branch of flavonoid biosynthesis. Time-resolved expression analysis with plants overexpressing the *KIN10* subunit of SnRK1 suggests a two-step mechanism that represses *MYBL2*, most likely involving HY5 during early time points and suppression of SnRK1 for persistent activation of anthocyanin biosynthesis in the course of high-light acclimation. We also found that PAP1-dependent repression of *FLS1, MYB12, MYB11, and MYB111* contributes to suppression of the flavonol branch, most likely by favouring the redistribution of precursor towards the anthocyanin branch. The presented results also indicate that the regulation of pathway genes is independent of flavonoid biosynthesis intermediates and end products.

## Results and Discussion

### SnRK1 acts upstream of *PAP1* for high-light induction of anthocyanin biosynthesis pathway genes *in planta*

Overexpression of the catalytic SnRK1 subunit *KIN10* prevented the activation of *DFR, LDOX*, transcription factors and anthocyanin accumulation in high-light (Zirngibl et al., 2023). This indicated that SnRK1 has to be inactivated to allow full activation of anthocyanin biosynthesis, most likely by permitting the activation of central transcription factors, such as PAP1 (Fig. 1). To provide genetic evidence that *PAP1* regulation involving SnRK1 is causative for high-light- and sugar-dependent control of anthocyanin biosynthesis, double overexpression lines were obtained by introducing the dominant *pap1-D* mutant allele (Borevitz et al., 2000) into *KIN10ox* (hereafter *pap1-D KIN10ox*). The two wild-types (WT), single overexpression and two *pap1-D KIN10ox* lines were exposed to high-light and sampled after 8 and 24 h for mRNA and anthocyanin accumulation, respectively. The *pap1-D KIN10ox* individuals showed a comparable overexpression of *KIN10* as revealed by western blot and gene expression analysis (Fig. 1 A-B). Notably, before the high-light shift (t0), *PAP1* was markedly overexpressed by approximately 20-fold in *pap1-D* and *pap1-D KIN10ox* compared to the respective WT (Fig. 1C). After 8 h high-light exposure, *PAP1* mRNA abundance increased by approximately 12-fold in Col-0 and Ler compared to t0. In contrast, in *pap1-D* mutant backgrounds, *PAP1* was increased by only 0.25-0.4-fold relative to t(0) (Fig. 1C). Hence, due to the marked induction in WT but only marginal increase in *pap1-D* and *pap1-D KIN10ox* upon high-light exposure, a comparable level of *PAP1* was detected in the genotypes. However, *PAP1* expression was lower in *KIN10ox* compared to both WT but fully recovered to *pap1-D* level in *pap1-D KIN10ox*.

**Figure 1:**
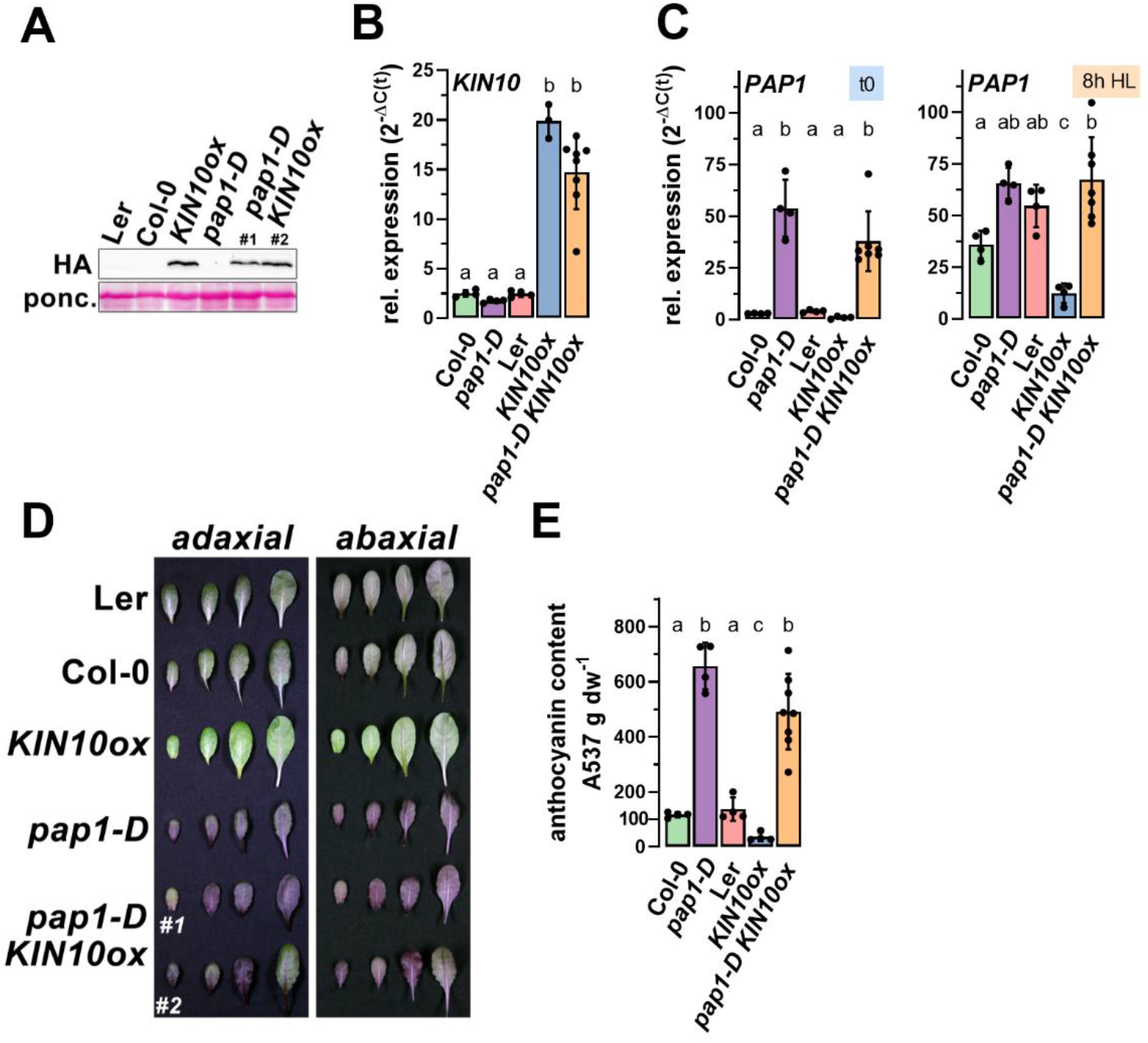
Overexpression of *PAP1* rescues diminished anthocyanin accumulation in *KIN10* overexpression lines. (**A**) Western blot detection of KIN10-HA protein in the *KIN10ox* overexpression lines using an HA-tag specific antibody. (**B and C**) Expression of *KIN10* (*SNRK1α1*) after 8 h HL and *PAP1* (*PRODUCTION OF ANTHOCYANIN PIGMENT1*) before (left) and after 8 h (right) of high-light exposure. (**D and E**) Photographs of representative rosette leaves (D) and anthocyanin content (E) in the indicated genotypes after 24 h high-light exposure. Relative gene expression is expressed as 2^-ΔC(t)^ using *SAND* as reference gene. The graphs show the mean of values (n≥4). Letters indicate significance groups identified with one-way ANOVA analysis (Tukey’s multiple comparison tests, adj p < 0.05). For *pap1-D KIN10ox* results of individuals from two independent progenies of crosses between *pap1-D* and *KIN10ox* (#1 and #2) are shown as one bar.

The WT leaves showed an intense purple colouration after high-light exposure (Fig. 1D). In contrast, *KIN10ox* leaves were much paler after the treatment (Fig. 1D), which was confirmed to be caused by a strongly diminished anthocyanin content in *KIN10ox* compared to WT (ratio *KIN10ox*/WT=0.25, Fig. 1E). Overexpression of *PAP1* resulted in a more pronounced purple colouration and a 6-fold increase of the anthocyanin content in *pap1-D* compared to WT (Fig. 1D-E). In contrast to *KIN10ox*, the *pap1-D KIN10ox* double mutant anthocyanin contents were as high as in the *pap1-D* single overexpression line (Fig. 1D-E).

Except for the bHLH transcription factor encoding gene *GLABRA3 (GL3*, Fig. 2A), the representative anthocyanin biosynthesis genes *DFR and LDOX*, and the bHLH transcription factor encoding gene *TT8* were expressed to similar levels in *pap1-D KIN10ox* and *pap1-D*, which was approximately two-to-four times the level detected in the respective WT backgrounds (Fig. 2A-B). Therefore, bypassing the SnRK1-dependent repression by overexpressing *PAP1* in *KIN10ox* fully rescued the deficient activation of *DFR* and *LDOX* and anthocyanin accumulation in high-light by stimulating the expression of MBW complex components (Figs. 1-2). *5MAT* and *SCPL10*, essential for the ‘decoration’ of the core anthocyanidin scaffold, were induced in *pap1-D* but repressed in *KIN10ox* and *pap1-D KIN10ox*, indicating that overexpression of KIN10 is epistatic to PAP1 overexpression (Fig. 2B).

**Figure 2:**
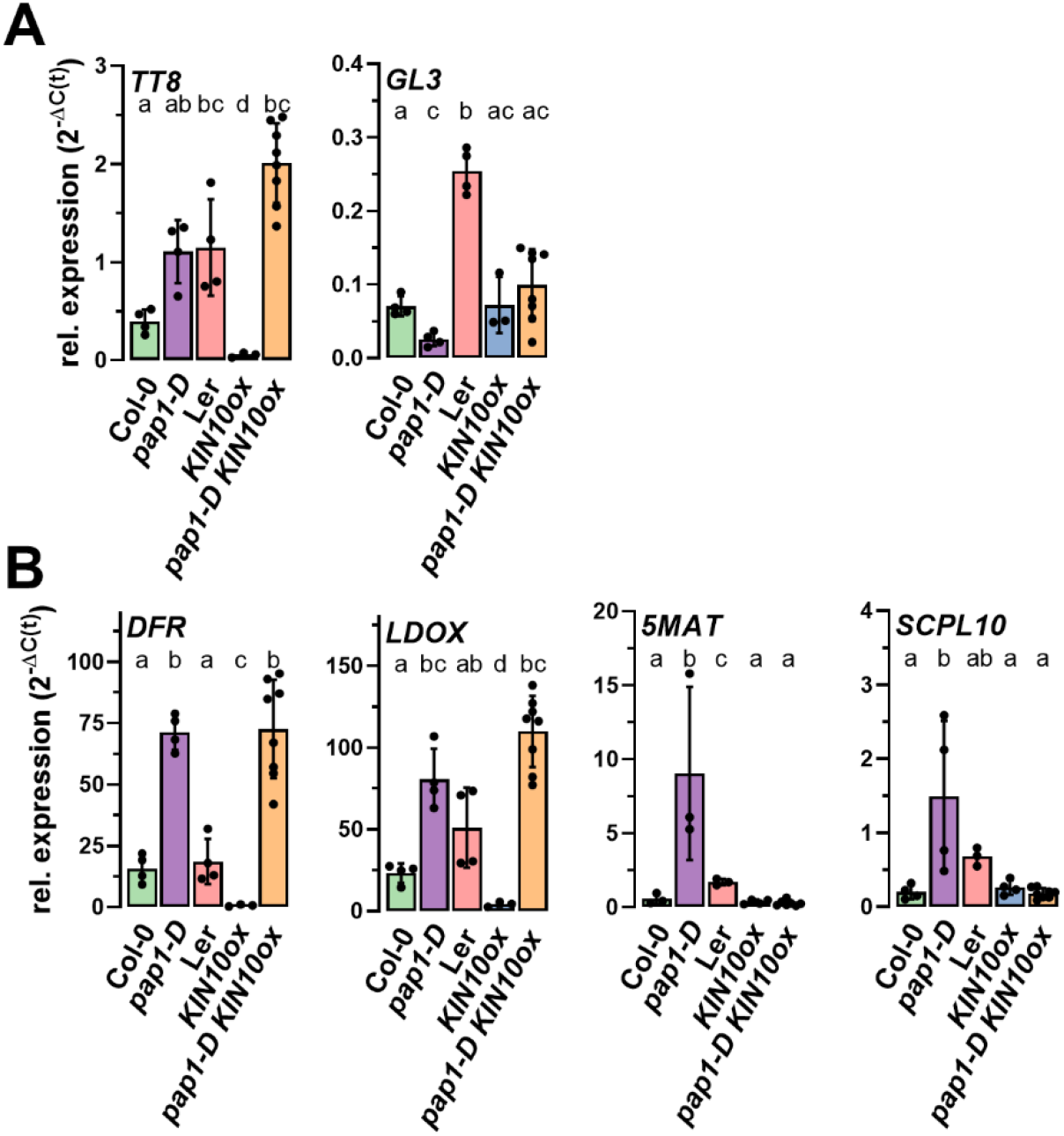
Repression of anthocyanin biosynthesis genes by *KIN10* overexpression is rescued by overexpression of *PAP1*. (A) Expression of *TRANSPARENT TESTA8 (TT8)*, and *GLABRA3 (GL3)* after 8h high-light treatment. (B) Expression of *DIHYDROFLAVONOL REDUCTASE (DFR)* and *LEUCOCYANIDIN DIOXYGENASE (LDOX*), *ANTHOCYANIN 5-O-GLUCOSIDE MALONYLTRANSFERASE* (*5MAT*), and *SINAPOYL-GLUCOSE TRANSFERASE* (*SCPL10*) after 8h high-light treatment. Relative gene expression is expressed as 2^-ΔC(t)^ using *SAND* as reference gene. The graphs show the mean of values ±SD (n≥4). Letters indicate significance groups identified with one-way ANOVA analysis (Tukey’s multiple comparison tests, adj p < 0.05).

Targeted gene expression analysis showed *MYB11, MYB12, MYB111, CHS*, and *CHI* repression in *KIN10ox* compared to WT (Fig. 3A-B). Also, *FLS1*, the key gene of the flavonol branch, was repressed in *KIN10ox* compared to WT in high-light (Fig. 3B). These results revealed a pronounced effect of SnRK1 on the expression of enzymes necessary for precursor production for all flavonoids and enzymes of the flavonol branch. While expression of *CHS* and *CHI* was rescued to at least WT-level, *FLS1, MYB11, MYB12*, and *MYB111* were still repressed in *pap1-D KIN10ox* compared to WT and expressed to similar levels as in *KIN10ox* (Fig. 3A-B). We also noted that strong expression of *PAP1* coincides with the pronounced repression of *FLS1* and *MYB11, MYB12, and MYB111* in both *pap1-D* mutant backgrounds compared to WT (Fig. 3A). Rhamnosyltransferase *UGT78D1* transferase expression was WT-like in all analysed genotypes. In contrast, the level of *OMT1* transcripts was induced in *pap1-D* but markedly repressed in *KIN10ox* mutant backgrounds (Fig. 3B).

**Figure 3:**
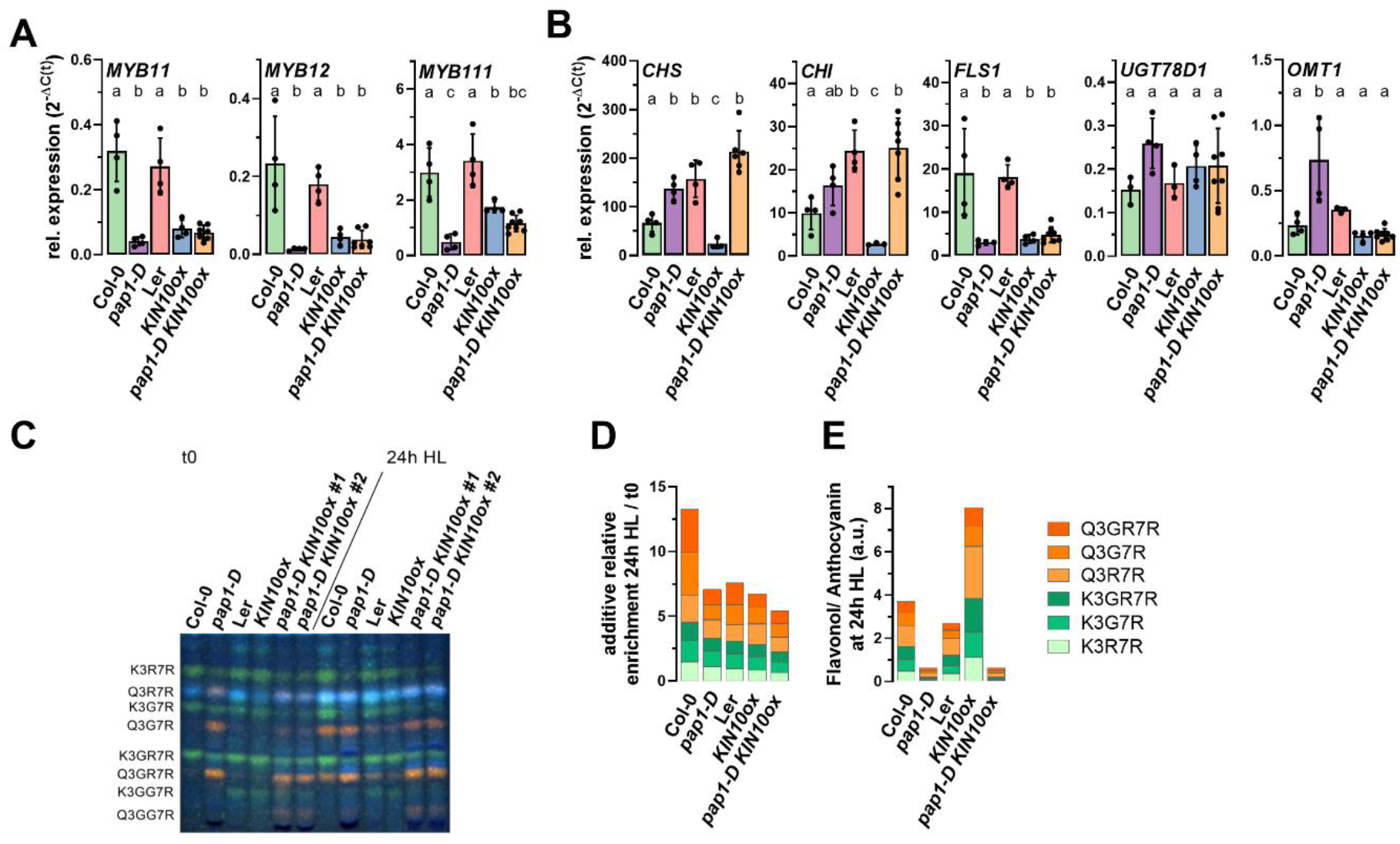
Ectopic expression of *PAP1* and *KIN10* affects the expression of general flavonoid biosynthetic genes and the flavonol branch. **(A)** Expression of *MYB11, MYB12*, and *MYB111* after 8h high-light treatment. **(B)** Expression of *CHALCONE SYNTHASE (CHS), CHALCONE ISOMERASE (CHI)*, and *FLAVONOL SYNTHASE1 (FLS1), FLAVONOL 3-O-RHAMNOSYLTRANSFERASE* (*UGT78D1*), and *FLAVONOL 3’O-METHYLTRANSFERASE1* (*OMT1*) after 8h high-light treatment. **(C)** HPTLC analysis of flavonol glycosides in plants before (t0) and after 24 h high-light exposure (24 h HL). Green, kaempferol derivative; orange, quercetin derivative; faint blue, sinapate derivative. K3R7R, kaempferol-3-O-rhamnoside-7-O-rhamnoside; Q3R7R, quercentin-3-O-rhamnoside-7-O-rhamnoside; K3G7R, kaempferol-3-O-glucoside-7-O-rhamnoside; Q3G7R, quercetin-3-O-glucoside-7-O-rhamnoside; K3GR7R, kaempferol 3-O-[rhamnosyl-glucoside]-7-O-rhamnoside; Q3GR7R, quercetin 3-O-[rhamnosyl-glucoside]-7-O-rhamnoside. Flavonols extracted from a pool of 24 rosette leaves from four plants and corresponding to 0.2 mg dry weight were separated in each lane. **(D)** Densitometric analysis and semi-quantitative comparison of pixel intensities evaluated from the HPTLC shown in (C). The ratio of the maximum peak heights/pixel intensities for each band before (t0) and after 24 h HL are shown. For *pap1-D KIN10ox* the mean value of line #1 and #2 is shown. **(E)** Ratio of the flavonol (maximum pixel intensity) and anthocyanin content (A537 g dw^-1^) after 24 h HL exposure. Values for the anthocyanin contents were obtained from Figure 1E. For *pap1-D KIN10ox* the mean value of line #1 and #2 is shown. Relative gene expression is expressed as 2^-ΔC(t)^ using *SAND* as reference gene. The graphs show the mean of values ±SD (n≥4). Letters indicate significance groups identified with one-way ANOVA analysis (Tukey’s multiple comparison tests, adj p < 0.05).

To test for a direct PAP1 function in the repression of *FLS1* and *MYB11, MYB12, MYB111* a *PAP1* knockout mutant (*pap1*) and the corresponding WT background (Nössen, Nos) were exposed to high-light (Fig. 4A). The time-resolved expression analysis showed a significant reduction of *CHS* and *FLS1* and a minor, though not significant, reduction of *MYB11, MYB12, MYB111* transcript levels after 8 h high-light exposure. The induction and repression of TT8 and MYBL2, respectively, were overall WT-like in *pap1* (Fig. 4B). In contrast, *DFR* and *LDOX*, two genes essential for anthocyanin production, were strongly repressed in the absence of *PAP1*, corroborating previous findings (Fig. 4B, Tohge et al., 2005, Zheng et al., 2019).

**Figure 4:**
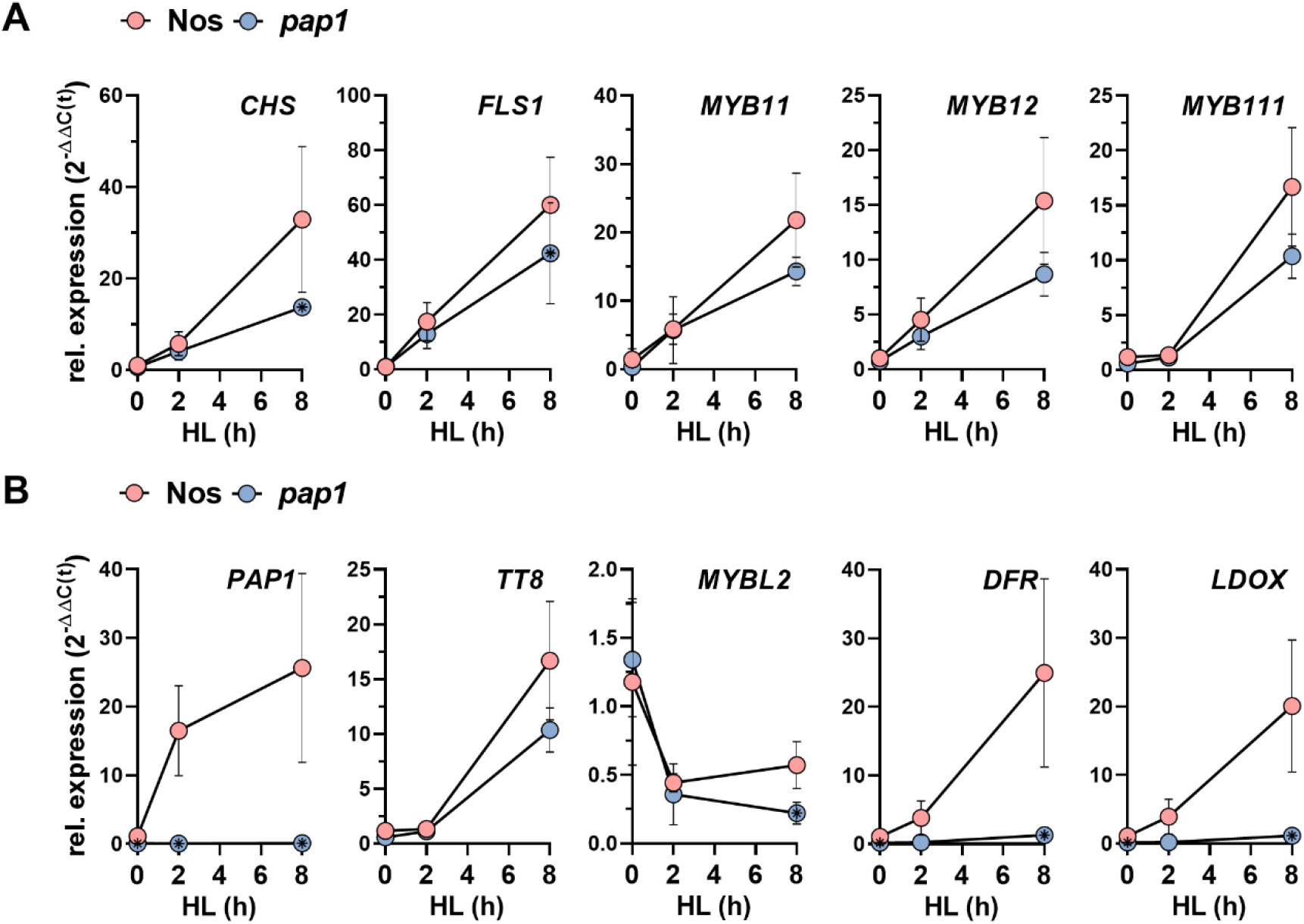
Gene expression analysis in *PAP1* knock out mutant (*pap1*) **(A)** Expression of *CHS, FLS1, MYB11, MYB12* and *MYB111* in the course of a high-light treatment. **(B)** Expression of *PAP1, TT8, MYBL2, DFR and LDOX* in the course of a high-light treatment. Samples were taken before, 2 h and 8 h after shifting the plant to high-light conditions. Nos = Nossen WT control for the *PAP1* knockout mutant *pap1*. Relative gene expression is expressed as 2^-ΔΔC(t)^ using *SAND* as reference gene and Nos before the high-light shift as control. The graphs show the mean of values ±SD (n≥3). Asterisks inside circles indicate significantly different values between Nos and *pap1* (p<0.05, student’s t-test).

Semi-quantitative analysis of flavonol glucoside accumulation revealed a pronounced accumulation of quercetin-type (orange in Fig. 3C) and a lower content of kaempferol-type (green in Fig. 3C) flavonol glycosides in *pap1-D* compared to WT before the HL shift (Fig. 3C, t0). While all flavonols were substantially increased in Col-0, only minor changes were detected in *pap1-D* after 24 h high-light exposure. The relative enrichment of quercetin- and kaempferol-type flavonol glycosides was reduced by 50% in *pap1-D* compared to Col-0 in high-light (Fig. 3D). Overexpression of *KIN10* in Ler background resulted in a 10% reduction of the relative flavonol glycosides enrichment in high-light. Compared to Col-0, diminished contents of kaempferol-type flavonol glycosides were observed in *pap1-D* before and after the high-light treatment (Fig. 3C). The *pap1-D KIN10ox* genotype exhibited lower levels of all flavonol glycosides compared to *pap1-D* single overexpression plants. The relative accumulation of flavonol glycosides within 24 h high-light was also reduced in the *pap1-D KIN10ox* genotype (Fig. 3D).

We next calculated the quotient of the anthocyanin content (Fig. 1E) and the cumulative maximum pixel intensities for the different flavonol derivatives obtained from the thin-layer chromatography (Fig. 3C) after 24 h high-light. This quotient does not represent the absolute distribution of precursors within the pathway but serves as a proxy for the relative distribution of precursors to the flavonol and anthocyanin branches. The parameter instead reflects the activation of the two branches relative to each other in the different genotypes. While Col-0 and Ler showed a similar flavonol/anthocyanin ratio, a more pronounced accumulation of anthocyanins relative to the flavonols was found in *pap1-D* (Fig. 3E). In contrast and besides an overall reduction in flavonol glycoside accumulation during high-light (Fig. 3D), overexpression of *KIN10* resulted in a marked increase of the flavonol/anthocyanin ratio compared to both WT backgrounds (Fig. 3E). This finding indicated a prevalent enrichment of flavonols compared to the anthocyanins in *KIN10ox*. Simultaneous overexpression of *KIN10* and *PAP1* diminished the flavonol/anthocyanin ratio to the same level observed for the *pap1-D* single overexpression line, suggesting that *PAP1* overexpression is epistatic to *KIN10ox* through a more pronounced channelling of intermediates towards the anthocyanin branch.

In summary, PAP1 and KIN10 overexpression negatively affected *MYB11, MYB12, MYB111* and *FLS1* expression. Compared to the single mutants, the lack of additive effects in the repression of these genes in the *pap1-D KIN10ox* would justify the assumption that PAP1 and KIN10 act in the same (signalling) pathway. Given the marked repression of *PAP1* by *KIN10* overexpression (Fig. 1C) and lack of induction in *pap1-D* it, however, seems unlikely that PAP1 is involved in SnRK1-dependent downregulation of *FLS1, MYB11, MYB12*, and *MYB111*. Additionally, WT-like expression of *KIN10* was observed in the *pap1-D* single mutant (Fig. 1B); thus, repression of *FLS1, MYB11, MYB12*, and *MYB111* is most likely independent of KIN10 (expression). MYB12 plays a role in leaf flavonol production and stimulates reporter gene expression driven by the *FLS1* promoter in transactivation assays (Mehrtens et al., 2005, Stracke et al., 2007). Based on an expression analysis conducted previously, strong overexpression of *MYB12* did not rescue *FLS1* expression in leaves of *pap1-D* (Nakabayashi et al., 2014). Hence, either the sole overexpression of MYB12 is insufficient, or MYB12 activity depends on (an) additional factor(s), in which case PAP1 overexpression is epistatic. Alternatively, PAP1 may act as a repressor of *FLS1* independent of the activating SG7 R2R3-MYB factors but possibly also involving other factors, such as R3-MYBs acting as repressors (LaFountain and Yuan, 2021). However, an impact of PAP1 either as a direct repressor or as a repressor of an activating factor on *MYB11, MYB12, MYB111, CHS*, and *FLS1* seems unlikely since the *pap1* knockout mutant did not show an overaccumulation but rather WT-like induction of these transcripts in standard and high-light conditions (Fig. 4). Notably, overexpression of *PAP1* resulted in suppression of *MYB11, MYB12, MYB111* and *FLS1* and lower flavonol glycoside levels during cold acclimation, indicating that fine-tuning of flavonoid biosynthesis via PAP1 is also relevant for other acclimation-relevant conditions (Schulz et al., 2016).

Our study provides genetic evidence for the importance of SnRK1 on *PAP1* expression *in planta* under acclimation-relevant conditions, which is supported by previous findings obtained with transient protoplasts assays and experiments involving exogenous sugar feeding (for example Baena-Gonzalez et al., 2007, Ramon et al., 2019). We propose that SnRK1 acts on *PAP1* by a yet unknown mechanism to prevent activation of flavonoid biosynthesis when it is not essential, for example, under normal light conditions, to avoid futile consumption of carbon skeletons and carbohydrates. On the other hand, the inactivation of SnRK1 by increasing sugar contents permits the activation of flavonoid biosynthesis for the production of photoprotective flavonoids in high-light. Our results suggest that SnRK1 predominantly regulates the anthocyanin branch of flavonoid biosynthesis (Fig. 3), which is conceivable considering the higher carbon demand for anthocyanin production compared to, for example, flavonols (Saito et al., 2013). This is further supported by the finding that SnRK1 dominantly represses also gene products required for the final steps of anthocyanin production, such as SCPL10 for anthocyanin glycosylation (Fig. 2). In Arabidopsis, eleven anthocyanin derivatives with different degrees of ‘decoration’ are formed (Saito et al., 2013), all of which absorb green-yellow light in acidic conditions and hence give a purple/pink appearance. The anthocyanin quantification method used here is based on acidic extraction and photometric detection at 537 nm (green-yellow), which does not discriminate between the different anthocyanins. Hence, we believe that *pap1-D KIN10ox* plants, although recovered to *pap1-D*-like level due to the repressed 5MAT and SCPL10 (Fig. 2B), most likely do not accumulate the identical spectra of anthocyanin derivatives as *pap1-D* single overexpression lines. An HPLC(/MS) approach should be employed to analyse these differences in more detail.

Repression of *PAP1* is independent of the other subunits of the SnRK1 complex (Ramon et al., 2019). Therefore, a direct impact of the catalytic α subunit KIN10 on *PAP1* induction could be assumed. Although the involved factor(s) downstream of SnRK1 and upstream of *PAP1* remain to be identified, the dominant transcriptional regulation of *PAP1* by SnRK1 acts in concert with the post-translational regulation of the MBW components (Broucke et al., 2023) and pathway enzymes (Wang et al., 2021), ensuring the sharp regulation of energy- and carbon-demanding flavonoid biosynthesis in fluctuating environmental conditions.

### A two-step mechanism involving SnRK1 affects *MYBL2* repression in high-light

In the course of expression analysis of important factors affecting anthocyanin accumulation, we found that *MYBL2*, a repressor of MBW complex activity (Matsui et al., 2008), was repressed shortly after the transfer to high-light and maintained a low expression level in WT during the following period of the treatment (Fig. 5A). In contrast, the *KIN10ox* lines showed a two-phasic *MYBL2* expression. Within the first hours of high-light exposure, a WT-like repression of *MYBL2* was observed for the *KIN10ox* lines, whereas after four hours, *MYBL2* increased again to the level before the high-light shift in *KIN10ox* (Fig. 5A). While corroborating the induction of *MYBL2* by *KIN10* overexpression in protoplast assays (Baena-Gonzalez et al., 2007, Broucke et al., 2023) our results suggest a previously unrecognised two-step mechanism for the sustained repression of *MYBL2*, which was not the case in the later phase of high-light exposure when *KIN10* was overexpressed (Fig. 5). Given that MYBL2 diminishes MBW complex activity through a concurrent interaction with PAP1 (Dubos et al., 2008, Matsui et al., 2008), more pronounced expression of *MYBL2* contributes to the diminished expression of flavonoid biosynthesis genes and anthocyanin accumulation in *KIN10ox*. However, further analysis is required to provide experimental evidence that MYBL2 acts downstream of SnRK1.

**Figure 5:**
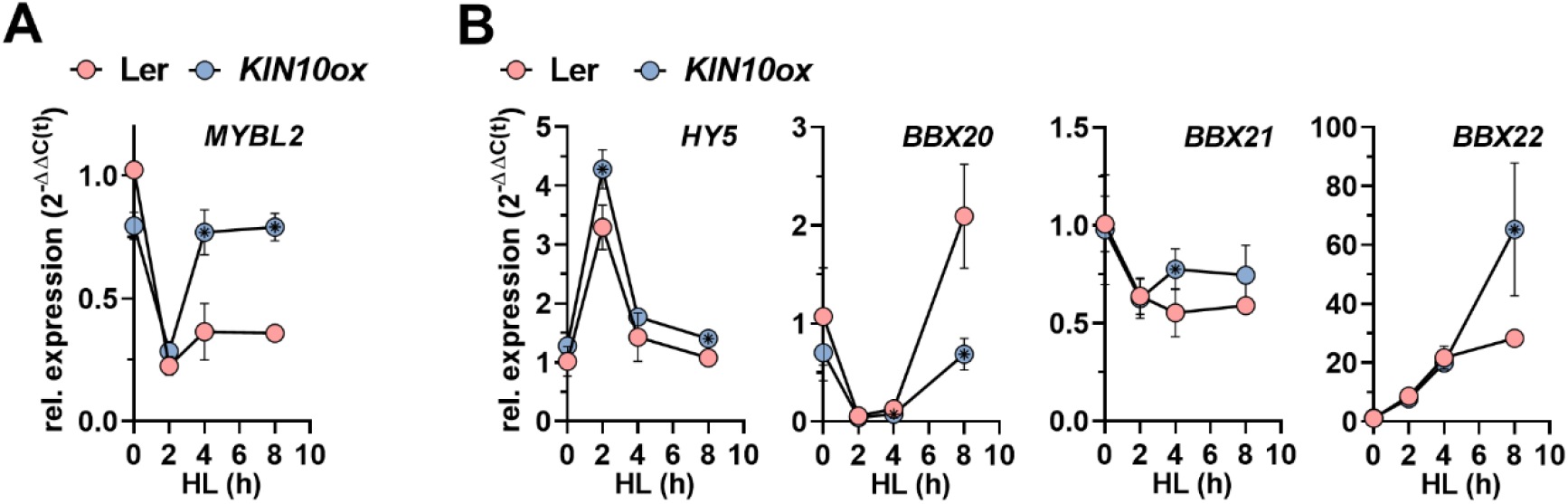
Expression analysis of *MYBL2, HY5* and *BBX20, BBX21, BBX22* in the course of high-light acclimation. **(A)** Expression of *MYBL2* and (**B**) *HY5, BBX20, BBX21, BBX22* in Ler and *KIN10ox* in the course of high-light acclimation. Relative gene expression in relation to WT (Ler) before the high-light shift (t0) expressed as 2^-ΔΔC(t)^ using *SAND* as reference gene. The graphs show the mean of values ±SD (n≥4).. Asterisks indicate significantly different values between Ler and *KIN10ox* (p<0.05, student’s t-test).

Another transcription factor, HY5, is known to regulate flavonoid biosynthesis pathway genes and functions as an activator and repressor of *PAP1* and *MYBL2*, respectively (Shin et al., 2013, Wang et al., 2016). Although significantly increased at two time points, *HY5* showed a WT-like expression trajectory in the course of the high-light shift experiment in *KIN10ox* (Fig. 5B). Within two hours, HY5 expression reached a maximum and declined again after four and eight hours of high-light exposure. Without excluding changes at the post-translational level, these changes in *HY5* abundance make it unlikely that they are causative for the reduced *PAP1* activation in *KIN10ox* (Fig. 1C). We also suggest that the immediate induction of *HY5* (Fig. 5B) mediates the observed repression of *MYBL2* (Fig. 5A) in WT and *KIN10ox* shortly after the onset of high-light. This assumption is consistent with the previously reported repression of *MYBL2* by HY5 (Wang et al., 2016).

HY5 lacks a trans-activation domain and depends on cofactors, such as BBX20, BBX21, and BBX22, to function in photomorphogenesis and flavonoid biosynthesis regulation (Oyama et al., 1997, Bursch et al., 2020). Hence, we also tested WT and *KIN10ox* for the expression of these HY5 cofactors. *BBX20* expression decreased within the first four hours but then increased strongly until eight hours in WT and *KIN10ox*. However, the late induction of *BBX20* was significantly diminished in *KIN10ox* (Fig. 5B). Although some minor differences were observed, *BBX21* expression was comparable between the two genotypes. *BBX22* mRNA showed a continuous increase upon high-light shift and was more strongly enriched in *KIN10ox* compared to WT eight hours after the transfer to high-light. Although HY5 must also depend on other cofactors and there is functional redundancy between BBX20, BBX21, and BBX22 (Bursch et al., 2020), suppression of *BBX20* is likely to contribute to the downregulation of pathway genes in *KIN10ox*. Also, a more pronounced activation of *BBX22* could compensate for the repression of *BBX20*.

In summary, we propose that two regulatory mechanisms underpin the activation of anthocyanin biosynthesis. Previous results (Wang et al., 2016) and our gene expression analysis suggest that HY5 may act on the pathway in the short term by repressing *MYBL2*. In the long-term, repression of SnRK1 by rising (regulatory) sugars permits the sustained activation of the pathway in high-light conditions (Zirngibl et al., 2023). This assumption is supported by the finding that *PAP1* expression initially increased but then decreased in a triosephosphate/phosphate translocator (*tpt-2*) mutant with deficient sugar- and SnRK1 signalling and activation of the anthocyanin branch of flavonoid biosynthesis in high-light (Zirngibl et al., 2023). Since *hy5* shows diminished but still accumulates substantial anthocyanin accumulation under high-light (Kleine et al., 2007), HY5 is essential but insufficient to fully activate anthocyanin biosynthesis in high-light. As it exerts a positive and negative function on other transcription factors and regulators to balance anabolic and catabolic reactions (summarised in Li et al., 2021), it is conceivable that SnRK1 also governs the control of energy- and carbohydrate-consuming anthocyanin biosynthesis. Although transcript analysis may exclude an impact of SnRK1 on *HY5*, we do not exclude post-translational mechanisms regulating the expression and/or stability of HY5, its cofactors or MYBL2.

### Flavonoids do not regulate flavonoid biosynthesis pathway genes in high-light conditions

The repression of *MYB11, MYB12* and *MYB111* and their target *FLS1* in the anthocyanin-rich *pap1-D* mutants (Fig. 3A) raised the hypothesis that the more pronounced activation of the anthocyanin branch by PAP1 might be promoted by the suppression of other branches regulated by these MYB factors, such as for flavonols. We further speculated that an intermediate of the pathway or anthocyanins themselves exert a signalling function to regulate *MYB11, MYB12* and *MYB111*. Given that PAP1 most likely does not function as a direct repressor of the flavonol branch (Fig. 4), this assumption could explain the downregulation of *MYB11, MYB12, MYB111, CHS, FLS1* in anthocyanin-rich tissues (e.g., *pap1-D*).

To elucidate the potential function of flavonoids in the high-light-dependent regulation of flavonoid biosynthesis, a mutant allele for *CHALCONE SYNTHASE* (*CHS, tt4-15*) was introduced into the *pap1-D* background (Fig. 6A). The double mutant, was characterised by a WT-like phenotype but anthocyanin deficiency after high-light treatment (Fig. 6B-C). Compared to WT, overexpression of *PAP1* resulted in a marked stimulation of *TT8, DFR* and *LDOX* in *pap1-D* and *tt4 pap1-D* eight hours in the high-light (Fig. 6D). *CHS* deficiency did not result in any expression changes of the anthocyanin biosynthetic genes either without or with *PAP1* overexpression. Likewise, mRNA for *MYB11, MYB12* and *MYB111*, which regulate *CHS, CHI* and *FLS1*, were expressed at WT-like levels in *tt4* during high-light acclimation (Fig. 6E). We also found a *pap1-D*-like overexpression of *CHI* in *tt4 pap1-D*, suggesting that a PAP1-dependent mechanism acting early in the pathway independent of flavonoids in high-light exists (Fig. 6E). Notably, knockout of *CHS* in the background of *pap1-D* resulted in the same repression of *MYB11, MYB12*, and *MYB111* as observed in *pap1-D* during high-light acclimation (Fig. 6E). In contrast to *CHS, FLS1* was repressed in *tt4 pap1-D* to the same extent as in the *pap1-D* overexpression line (Fig. 6E).

**Figure 6:**
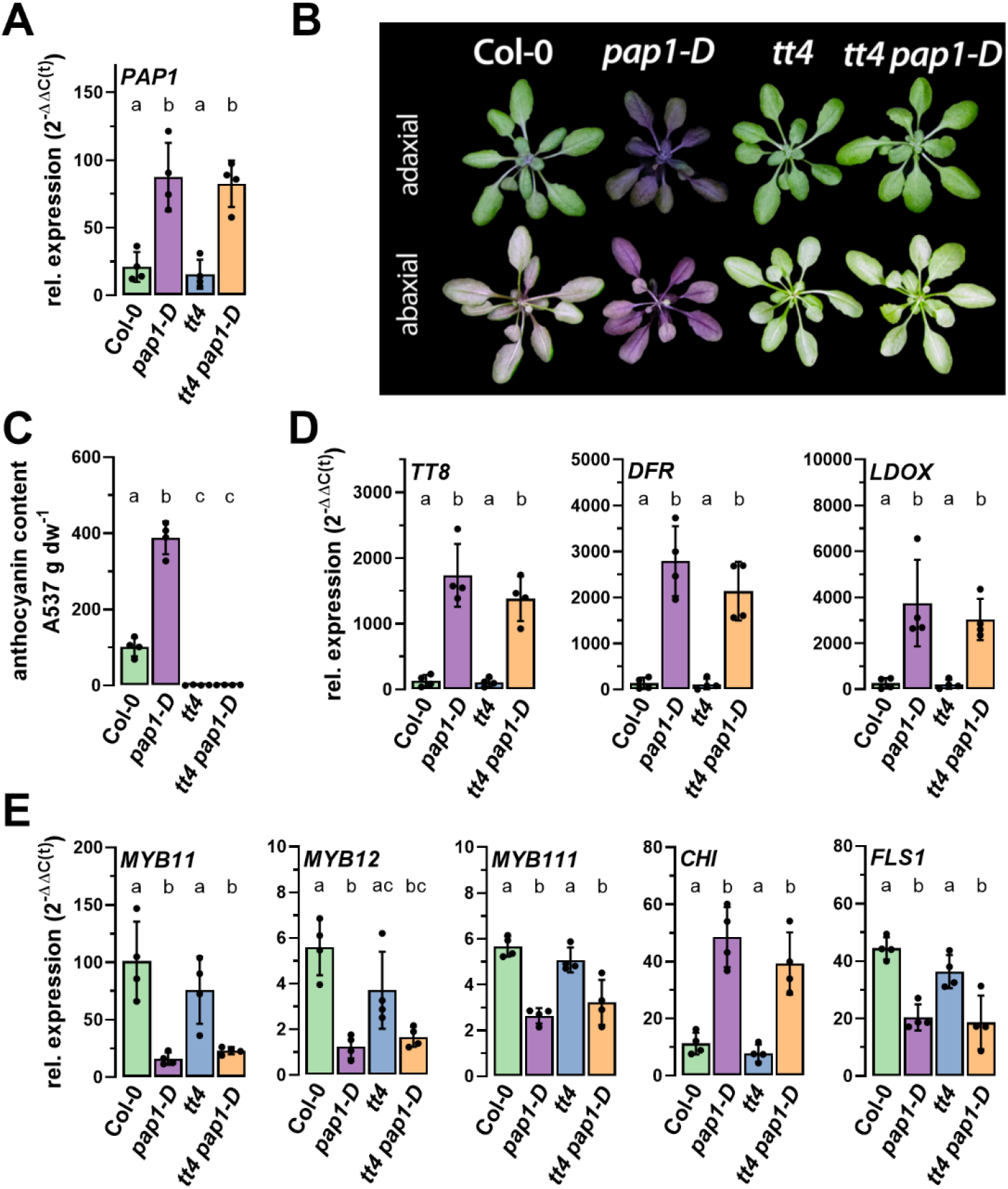
*PAP1* overexpression in a chalcone synthase (*tt4*) mutant background during high-light acclimation. **(A)** Expression of *PAP1* in the indicated genotypes. **(B)** Phenotype and (**C**) anthocyanin content in WT and mutant plants after 24 h high-light exposure. (**D**) Expression of *TT8, DFR* and *LDOX* and (**E**) *MYB11, MYB12, MYB111, CHI* and *FLS1* after 8 h high-light exposure. Relative gene expression in relation to Col-0 before the high-light shift expressed as 2^-ΔΔC(t)^ using *SAND* as reference gene. The graphs show the mean of values ±SD (n≥4). Letters indicate significance groups identified with one-way ANOVA analysis (Tukey’s multiple comparison tests, adj p < 0.01).

Taken together, we concluded that under our experimental conditions, intermediates or end-products of flavonoid biosynthesis do not mediate suppression of the flavonol branch or participate in the control of flavonoid biosynthesis in general. However, in other experimental conditions and set-ups, a function of flavonoids in biotic interactions in the rhizosphere (Hassan and Mathesius, 2012) and the circadian clock (Hildreth et al., 2022) have been reported. Although first insights have been gained (Nakabayashi et al., 2014), further experiments are needed to shed light on the signalling function of flavonoids or specific flavonoid derivatives in acclimation-relevant conditions. Changes in the light response originating from the massive accumulation of UV light-absorbing anthocyanins and internal shading, which might affect the (UV-) light-dependent accumulation of *MYB12* or *CHS* (Podolec et al., 2022) in *pap1-D*, seem unlikely given the absence of flavonoids but the repression of these transcripts in *tt4 pap1-D* (Fig. 6E).

The results presented here prove that the transcriptional repression of *MYB11, MYB12, MYB111* and *FLS1* indirectly results from PAP1 overexpression. As shown with the flavonoid-deficient *tt4 pap1-D* double mutant, a more pronounced accumulation of flavonoids is not causative for the suppression of the flavonol branch in anthocyanin-rich tissues (Fig. 6). It remains to be elucidated how *PAP1* overexpression contributes to the suppression of the flavonol branch at the transcriptional level. However, increased expression of structural anthocyanin biosynthesis genes resulted in a predominant channelling of precursors towards the anthocyanin branch (Fig. 3), fitting to the previously proposed pull- and push mechanisms of intermediates within the pathway (Jiang et al., 2020). This regulatory circuit is most likely responsible for the demand-driven distribution of intermediates between different flavonoid biosynthesis branches and the production of end products in high-light.

## Material and Methods

### Plant material and growth conditions

*Arabidopsis thaliana* (Arabidopsis) plants were grown on soil in short-day conditions (10 h light) at 100 µmol m^− 2^ s^− 1^ light at 22°C. As wild-type, Col-0 and Ler were used. The mutants *tt4-15, pap1-D* (Col-0 background), *KIN10ox* (Ler background), and *pap1* knockout (Nos background) were obtained from the Nottingham Arabidopsis Stock Centre (NASC). The *pap1-D KIN10ox* and *tt4-15 pap1-D* lines were obtained by crossing, and homozygous progenies were selected. For *pap1-D KIN10ox* two lines from independent crosses were selected. The *pap1-D KIN10ox* double overexpression line data was obtained with seed batches from independent mother plants selected after crossing (lines #1 and #2). The overexpression of *PAP1* was proven by the accumulation of anthocyanins under normal light (100 µmol m^− 2^ s^− 1^) and/or qPCR validation. Overexpression of HA-*KIN10* was verified by western blotting using anti-hemagglutinin (HA)-specific antibody.

For the high-light shift experiment, 5-6-week-old plants were shifted at the end of the night for 24 h to 500 µmol m^− 2^ s^− 1^ at 22°C in a Conviron Gen1000 equipped with LEDs. For the experiment with *pap1-D KIN10ox* lines and *tt4 pap1-D* lines, one sample consisted of 12 randomly sampled leaves from two plants, and at least three biological replicates were harvested. For the time-resolved expression analysis of Ler and *KIN10ox* lines, four plants grown and treated on different pots were combined in one sample and four biological replicates were analysed.

### Protein extraction and western blot

For the protein extraction, 200 µL of protein extraction buffer (56 mM Na_2_CO_3_, 56 mM dithiothreitol DTT, 2% (w/v) SDS, 12% (w/v) sucrose, 2 mM EDTA) were added to 2 mg of leaf powder and mixed. The samples were incubated for 10 min at room temperature, another 20 min at 70°C and then centrifuged (10 min, 13000 rpm, RT). Protein extracts were used for a 12%-SDS-PAGE and blotted onto a nitrocellulose membrane. Ponceau staining was used for loading control. The membrane was blocked for 1 h at RT in 4% milk solution in TBS-T (12.4 mM Tris, 150 mM NaCl, 0.1% (v/v) Tween-20, pH 7.4), washed and incubated overnight at 4°C in 1% milk solution in TBS (12.4 mM Tris, 150 mM NaCl, pH 7.4) with the primary rabbit anti-HA-tag (1:5000) antibody. On the following day, the membrane was washed and incubated for 2 h at RT in 1% milk solution in TBS with the secondary goat anti-rabbit IgG coupled with horseradish peroxidase (1:5000). The signal detection was performed using Clarity Western ECL substrate (Bio-Rad, Germany) and an ECL Chemostar imager (Intas, Germany).

### Anthocyanin content measurements

Anthocyanin quantification was performed following a previously published protocol (Lotkowska et al., 2015). 2 mg of leaf powder was mixed with 1 mL anthocyanin extraction buffer (1% (v/v) HCl, 18% (v/v) 1-propanol, in water) and incubated for 3 h at RT in dark and mixed regularly. Samples were centrifuged (15 min, 13000 rpm, RT) and transferred to cuvettes. The absorption was measured at 537, 650 and 720 nm, and the anthocyanin absorption was calculated with the following formula: ((A_537_-A_720_)-0.25x(A_650_-A_720_)) dw^-1^

### RNA extraction and qPCR analysis

RNA was extracted as previously described in Oñate-Sánchez and Vicente-Carbajosa (2008). 2 mg of leaf powder was mixed with 300 µL RNA extraction buffer (2% (w/v) SDS, 68 mM sodium citrate, 132 mM citric acid, 1 mM EDTA) and incubated for 5 min at RT. 100 µL DNA/protein precipitation solution (4 M NaCl, 16 mM sodium citrate, 32 mM citric acid) was added. Samples were mixed, incubated for 10 min on ice, and centrifuged for 10 min, 13000 rpm at 4°C. 350 µL of supernatant was transferred to a new tube and again centrifuged (10 min, 13000 rpm, 4°C). 300 µL of supernatant was mixed with 300 µL isopropanol, centrifuged (5 min, 13000 rpm, RT), and the supernatant was discarded. The pellet was washed with 800 µL 70% ethanol and centrifuged (5 min, 13000 rpm, RT). Ethanol was discarded, the pellet was dried, and then resuspended in 20 µL DEPC-H_2_O for 10 min on ice. 1.5 µg RNA were treated with DNAse I (Thermo Fisher) and transcribed to cDNA using RevertAid reverse transcriptase (Thermo Fisher) as described in the manufacturer’s protocol. For the qPCR, 1 µL of diluted cDNA (1:5) was used with 2 x ChamQ Universal SYBR qPCR Master Mix (Vazyme, China) and primers depicted in Table S1 in 6 µL-reactions. Samples were analysed with a CFX96-C1000 96-well plate thermocycler (Bio-Rad, Germany). Gene expression was quantified using either 2^-ΔC(t)^ or 2^-ΔΔC(t)^ method (Livak and Schmittgen, 2001) and was calculated and expressed relative to *SAND* (AT2G28390) as a reference gene. In the case of the *tt4 pap1-D* lines, expression was additionally normalised to the expression in WT before the high-light shift (t0; equals 1; 2^-ΔΔC(t)^). No technical replicates were analysed.

### High-performance thin layer chromatography (HPTLC) of flavonols

Flavonols were extracted from a pool of rosette leaves, which were homogenised in a mixer mill (Retsch, Haan, Germany) and freeze-dried. 20 µL of 80 % methanol was added per mg of sample dry weight and the samples were incubated at 60°C for 10 min on a thermoshaker. The samples were centrifuged at 20,000 g for 10 min. 4 µL of the resulting supernatants (corresponding to extract of 0.2 µg dry weight) were analysed by HPTLC on a Silica Gel 60 plate (Merck, Darmstadt, Germany) using a mobile phase consisting of ethyl acetate, formic acid, acetic acid and water (100:26:12:12) as described in (Stracke et al., 2007). Flavonol glycosides were detected by DPBA/PEG staining and UV light (Sheahan and Rechnitz, 1992).

## ACKNOWLEDGEMENTS

This project is supported by a grant from the German Research Foundation (DFG) in the framework of the TRR175, project C06 to ASR (INST 86/2042-1) and the BMBF.

## AUTHOR CONTRIBUTIONS

JD and RS performed the analysis, evaluated the data and contributed to the writing. ASR, conceived and supervised the study, evaluated the data, acquired funding and wrote the manuscript.

